# The indoor mycobiome of daycare centers is affected by occupancy and climate

**DOI:** 10.1101/2021.10.28.466379

**Authors:** Eva Lena F. Estensmo, Synnøve Smebye Botnen, Sundy Maurice, Pedro M. Martin-Sanchez, Luis Morgado, Ingeborg Bjorvand Engh, Klaus Høiland, Inger Skrede, Håvard Kauserud

## Abstract

Many children spend considerable time in daycare centers and may here be influenced by indoor microorganisms, including fungi. In this study, we investigate the indoor mycobiome of 125 daycare centers distributed along strong environmental gradients throughout Norway. Dust samples were collected from doorframes outside and inside buildings using a citizen science sampling approach. Fungal communities in the dust samples were analyzed using DNA metabarcoding of the ITS2 region. We observed a marked difference between the outdoor and indoor mycobiomes. The indoor mycobiome included considerably more yeasts and molds compared to the outdoor samples, with *Saccharomyces, Mucor, Malassezia* and *Penicillium* among the most dominant fungal genera. Changes in the indoor fungal richness and composition correlated to numerous variables related to both outdoor and indoor conditions; there was a clear geographic structure in the indoor mycobiome composition that mirrored the outdoor climate, ranging from humid areas in western Norway to drier and colder areas in eastern Norway. Moreover, the number of children in the daycare centers, as well as various building features, influenced the indoor mycobiome composition. We conclude that the indoor mycobiome in Norwegian daycare centers is structured by multiple factors and is dominated by yeasts and molds. This study exemplifies how citizen science sampling enables DNA-based analyses of a high number of samples covering wide geographic areas.

**Importance:** With an alarming increase in chronic diseases like childhood asthma and allergies, there is an increased focus on the exposure of young children to indoor biological and chemical air pollutants. Our study of 125 daycares throughout Norway demonstrates that the indoor mycobiome not only reflects co-occurring outdoor fungi but includes a high abundance of yeast and mold fungi with an affinity for indoor environments. A multitude of factors influence the indoor mycobiome in daycares, including building type, inhabitants, as well as the outdoor environment. Many of the detected yeasts and molds are likely associated with the human body, where some have been coupled to allergies and respiratory problems. Our results call for further studies investigating the potential impact of the identified daycare-associated mycobiomes on children health.

## Introduction

Over a few thousand years, humans have moved from a mostly outdoor lifestyle to spending a large part of their life in indoor environments. Although the diversity of other co-occurring organisms is considerably lower in indoor environments, humans are not alone. If moisture and organic materials are available indoors, fungi can grow and disperse spores. Some of the most prevalent fungi that are able to grow and sporulate in houses include various ascomycete molds, such as *Cladosporium, Penicillium* and *Aspergillus* (1, 2). Fungal growth can lead to poor indoor air quality, and some of these fungi are associated with allergic reactions (3-5) and respiratory disease symptoms (6, 7), which may have long-term impacts on human health. Furthermore, certain combinations of indoor fungi and bacteria in moisture-damaged buildings may also cause negative health effects, even in low concentrations (8).

In many countries, children spend considerable time in daycare centers, where they are exposed to indoors microorganisms, including fungi. Since young children often vector organic material such as soil and litter from nature, daycare centers may accumulate extra organic substrates promoting fungal growth, as compared to other indoor environments. In line with this, it has previously been shown that the concentration of fungi in daycare centers is higher compared to homes (9). In several studies, the outdoor environment has been reported as the main source of indoor fungi (10-13) due to the influx of spores through windows, entrances and the ventilation system. Hence, the vegetation and climate that structure the outdoor fungi will indirectly also structure the indoor mycobiome (11). In correspondence with this, in a recent DNA metabarcoding study performed in 271 private homes across Norway, we showed that outdoor climate was one of the main drivers of the indoor dust mycobiome (13). A similar observation was done by Barberán et al. from dust samples collected on the outside surface of homes across the USA (11).

In addition to the outdoor environment, the inhabitants and their diverse activities, the presence of pets and plants, as well as various building features, may contribute and structure the indoor mycobiome (14, 15). Many yeasts, such as *Malassezia* and *Candida*, are associated with the human body and may therefore be prevalent indoors (16-19). Which fungi that are associated with the human body may, to some extent, be age-dependent. For instance, the basidiomycete yeast *Malassezia* seems particularly prevalent on adults (20), while children tend to have a more diverse skin-associated mycobiome, including genera like *Aspergillus, Epicoccum, Cladosporium, Cryptococcus* and *Phoma*, in addition to *Malassezia* (18).

The indoor mycobiome can be analyzed in different ways, including isolation and cultivation of fungi, microscopy and different molecular analyses. DNA metabarcoding, based on high-throughput sequencing of PCR amplified markers, is established as an effective approach to survey fungal communities (21). In buildings, DNA metabarcoding of dust samples, integrating spores and hyphal remains that have accumulated over time, has proven to be an effective mean for exploring the indoor mycobiome (10, 12, 22-24). However, it might be difficult to get access and obtain samples from a representative number of buildings. By providing detailed instructions, dust samples can alternatively be collected by the inhabitants themselves, from which DNA can be extracted and analyzed further (24, 25). This type of community-based research, where networks of non-professionals help to collect data as part of a research project, is regarded as citizen science (26-28). Sampling through citizen science is a powerful approach, where sample equipment can be sent out by post, returning hundreds or even thousands of samples covering large geographic areas.

The impact long-term exposure to indoor fungi can have on human health highlights the need to better characterize the indoor mycobiome from an early age. In this study, we aim to analyze the indoor mycobiome associated with daycare centers. We ask (1) which outdoor and indoor factors drive the daycare mycobiome, and (2) which fungal groups dominate in the daycare centers, as compared to outdoor samples. To address these research questions, we chose a citizen science approach, where daycare personnel collected dust samples according to our instructions. We obtained 572 samples from doorframes inside (bathroom and main room) and outside (main entrance) of 125 daycare centers throughout Norway (Fig. 1). Norway spans extensive gradients in climate and other environmental drivers, enabling us to evaluate the influence of the outdoor environment on the indoor mycobiome, in addition to building features and inhabitant characteristics. The obtained dust samples were analyzed by DNA metabarcoding of the rDNA Internal Transcribed Spacer 2 (ITS2) region.

**Figure 1.**
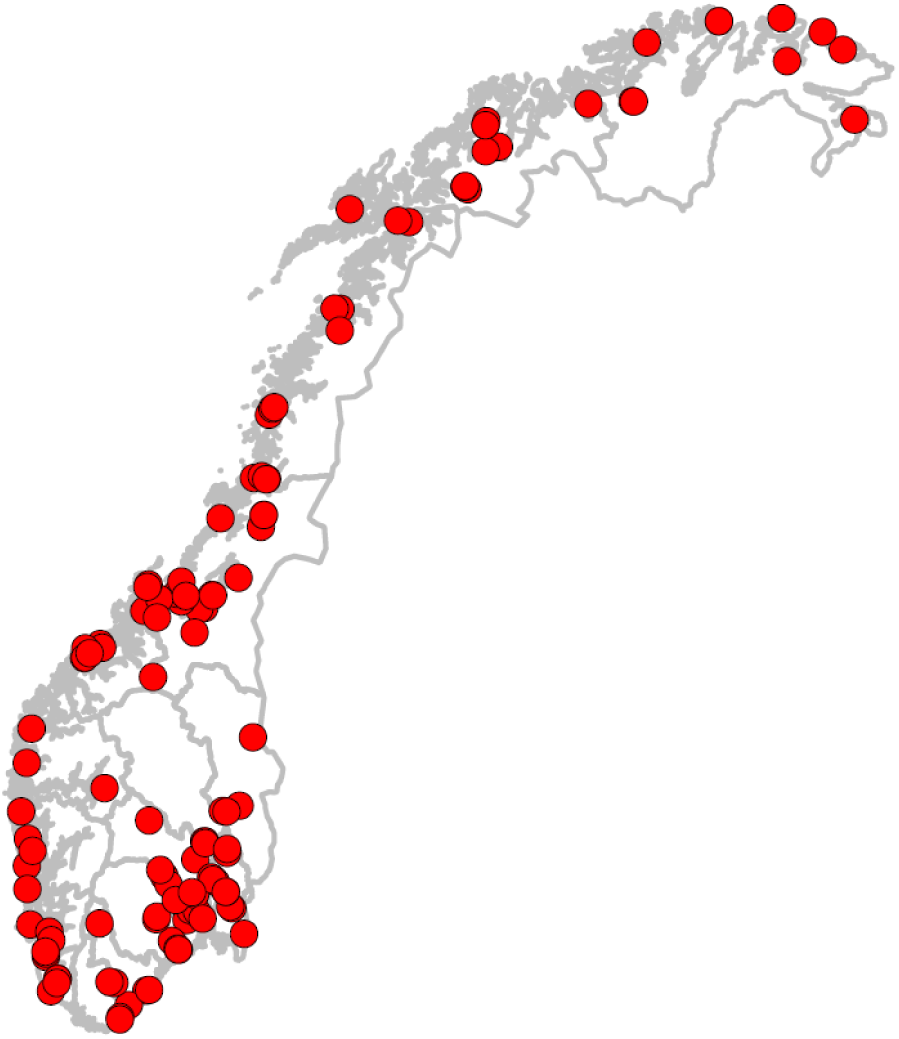
Map of Norway showing the geographical location of the 125 daycare centers included in the study. The samples were collected by citizen scientists, including samples from both indoor and outdoor environments.

## Results

### Factors influencing the indoor mycobiome

Our final rarified dataset from the 125 daycare centers included 748 836 sequences, with 1 342 sequences in each of the 558 samples from indoor and outdoor environments. A total of 5 946 fungal OTUs were detected in the dataset. In a multivariate (NMDS) analysis, we observed a relatively clear separation between the outdoor and indoor dust mycobiomes (Fig. 2a). However, the two types of indoor samples, main room versus bathroom, largely overlapped in fungal community composition (Fig. 2b). Through a questionnaire to the citizen scientists (personnel of the daycare centers), we obtained information about different building and occupancy variables (Table 1). In addition, information about the local climate and vegetation were extracted based on the geographic coordinates of the daycare centers (29). Considered individually, numerous of these variables correlated significantly with the compositional variation in the indoor mycobiome (Fig. 2c), including variables related to the daycare centers such as daycare type, construction year, number of departments, pests and building type. Climatic variables such as temperature and total insolation were also significantly correlated to the indoor mycobiome composition, as well as spatial variables that likely mirror additional regional environmental variability (Fig. 2c, d). Many of the inferred variables were associated with the major climate gradient stretching from humid, oceanic areas in western Norway, to inland, continental areas in eastern Norway (Fig. 2c, d). Evaluating the relative contribution of variables together in a CCA analysis (Table 2), revealed that longitude (mirroring the regional climate gradient), presence of pest/rodents, construction year of the daycare center and number of children were the main drivers of the fungal community composition, with very low interaction effects (<0.01%). However, these factors accounted altogether for only 7% of the variation in mycobiome composition (Table 2). The indoor fungal richness, calculated on a sample-basis, was significantly higher in the bathroom compared to the main room, and there was a significant positive correlation between indoor fungal richness and the maximum outside temperature during May at the sampling location, as well as the proximity to coast (see the Mixed Effect Model presented in Table 3).

**Figure 2.**
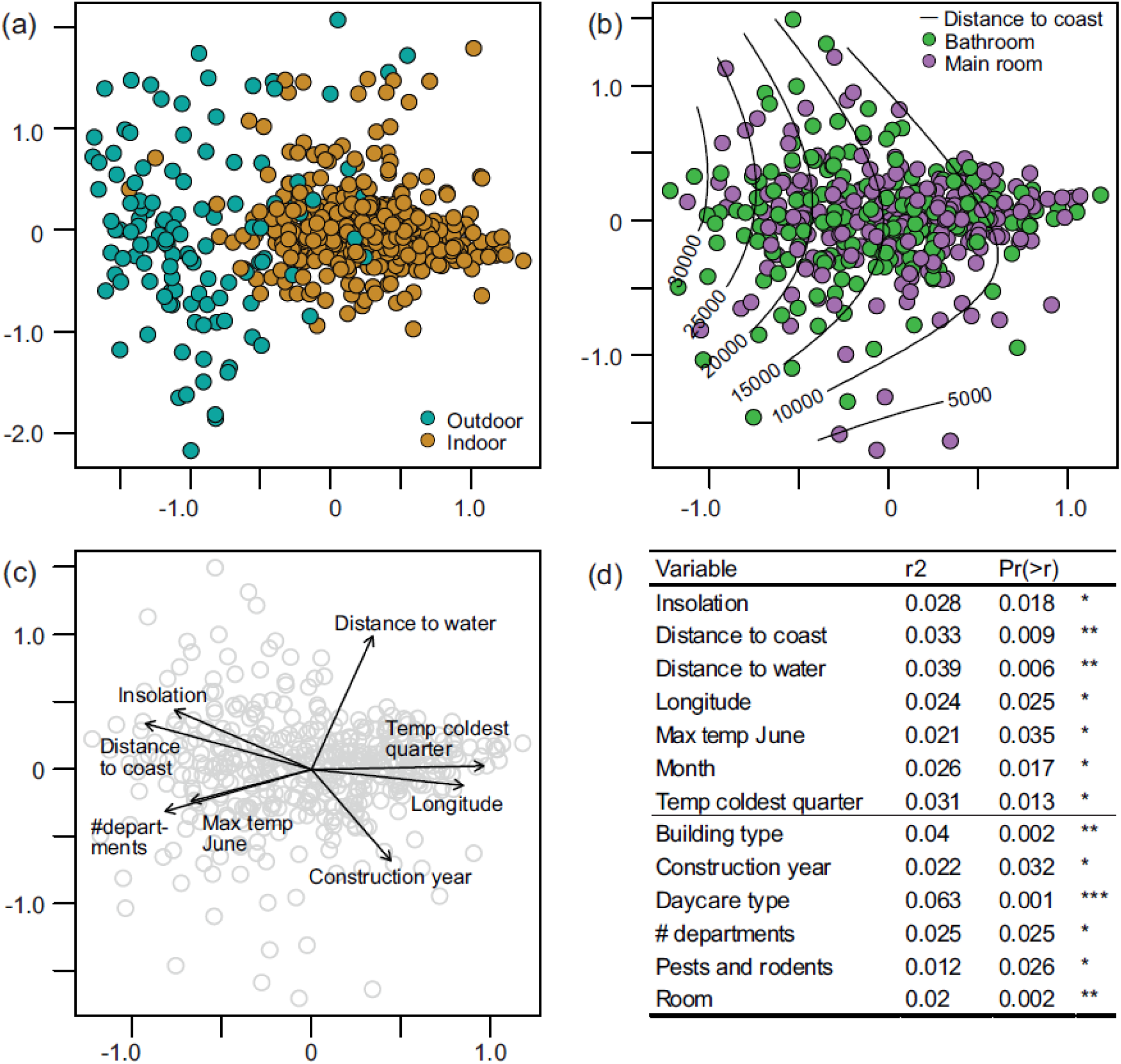
Fungal community composition in daycare centers. (a-c) Ordination plots displaying compositional variation in the dust mycobiome, where each point indicates one dust sample. (a) NMDS plot displaying both outdoor (cyan) and indoor (brown) samples. (b) NMDS plot of only indoor samples from bathrooms (green) and main rooms (purple). The isolines represent the distance to coast. (c) The indoor samples with vectors representing numeric variables with significant associations to the compositional variation in the indoor mycobiome (p<0.05). Categorical variables are not displayed. (d) Goodness-of-fit statistics (r2) for variables that significantly (p<0.05) account for variation in the composition of the indoor mycobiome. Variables related to regional climate are listed in the upper part of the table, while variables related to the specific daycares are listed in the lower part.

**Table 1.**
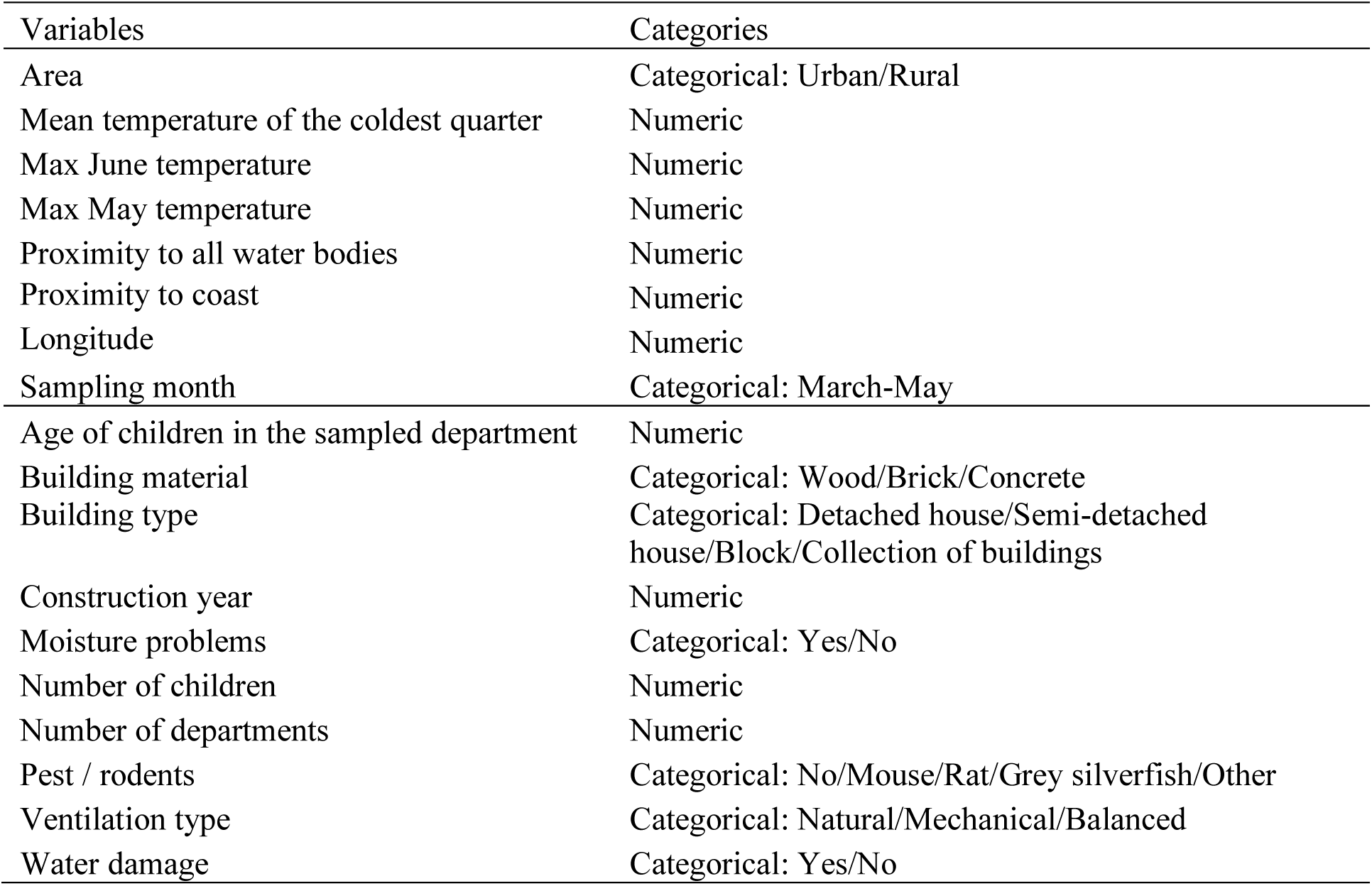
Climatic and building metadata selected by correlation test (|r| > 0.6). The upper part of the table include the six first climatic variables extracted from a database (29) using georeferences of the daycare centers. The variables about the occupants and building features provided by volunteers in each daycare center are listed in the lower part of the table.

**Table 2.** Variables with explanatory power in the Canonical correspondence analysis (CCA). Note that these variables may reflect a combination of variables or represent other variables not necessarily inferred here.

| Variables | Variation explained |
| --- | --- |
| Longitude | 0.0159 |
| Pests / rodents | 0.0187 |
| Construction year | 0.0181 |
| Number of children | 0.0156 |
| Interaction effects | 0.0001 |
| Unexplained variation | 0.9316 |

**Table 3.** Richness analyses using a mixed effect model with number of OTUs per sample as response and with daycare ID as a random effect. The variable Room type Bathroom is in the baseline of the model, the estimate for Room represents the difference from bathroom to main room.

| Variable | Estimate | Std error | t-value | p-value |
| --- | --- | --- | --- | --- |
| Room (bathroom = baseline) | -3.0773 | 1.310563 | -2.348083 | 0.0195 |
| Proximity to coast | 0.000095 | 0.000043 | 2.193266 | 0.0291 |
| Max May temperature | 1.671905 | 0.55933 | 2.98912 | 0.003 |

### Taxonomic composition of daycare mycobiome

In the indoor mycobiome, Saccharomycetales and Mucorales were most common, in contrast to the outdoor mycobiome that was mainly dominated by Pucciniales, Capnodiales, Agaricales and Chaetothyriales (Fig. 3a). The true yeasts of Saccharomycetales were considerably more abundant in the indoor environments. Malasseziales, a basidiomycete yeasts, was also somewhat more abundant indoors (Fig. 3a). We annotated the 1253 most abundant OTUs (OTUs with > 20 sequences) into different growth and life forms, which revealed that yeasts, dimorphic yeasts and molds were considerably more abundant in indoor environments, while saprotrophs, plant pathogens and lichens dominated relatively more in the outdoor samples (Fig. 3b).

**Figure 3.**
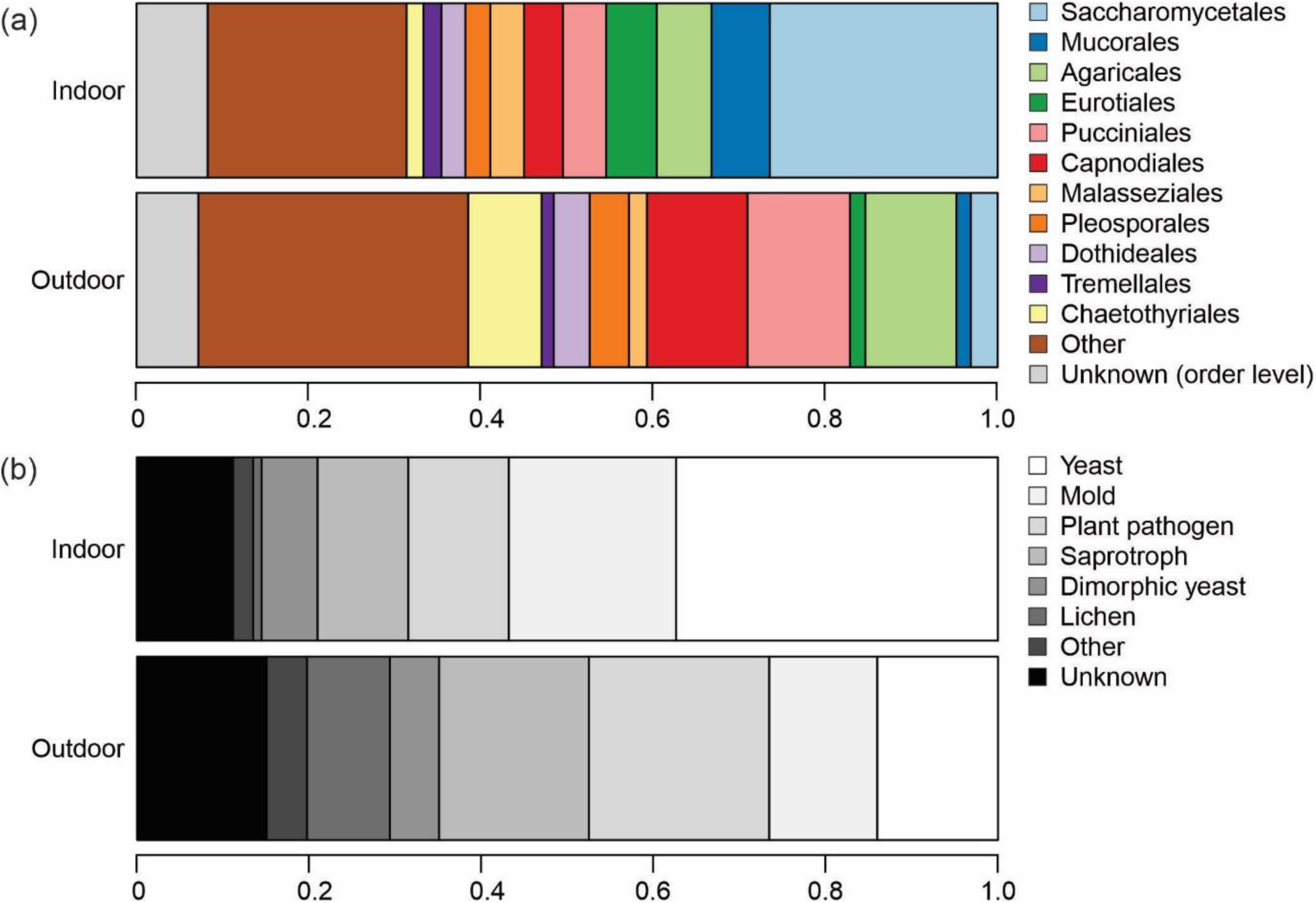
Taxonomic distribution of fungal OTUs in outdoor and indoor dust samples from the daycare centers reflecting sequence numbers. (a) Relative abundance of the main fungal orders based on the rarified OTU table. (b) Relative abundances of same OTUs annotated to different growth forms / nutritional modes. The category saprotrophs represents litter and wood decay fungi.

Among the top 30 genera detected in this study, measured in sequence abundance in a balanced indoor/outdoor dataset (where the sequence abundance of the two indoor samples were averaged), many had a clear affinity towards either indoor or outdoor environments (Fig. 4). Ten genera, namely, *Aspergillus, Candida, Debaryomyces, Filobasidium, Malassezia, Mortierella, Mucor, Penicillium, Rhodotorula, Saccharomyces* and *Wallemia*, had a clear affinity towards indoor environments. *Saccharomyces* was by far the most abundant genera in the indoor environment, with about 12.5 time’s higher abundance indoors compared to outdoors. In contrast, plant pathogens like *Melampsora, Puccinastrum* and *Melampsoridium* were relatively more common in the outdoor samples. Interestingly, some genera with affinity to the outdoor environment, like *Verrucocladosporium, Scoliciosporum* and *Sordaria* were almost exclusively present in the outdoor samples, while others, like *Cladosporium, Melampsoridium* and *Lycoperdon*, were also abundant in the indoor environment.

**Figure 4.**
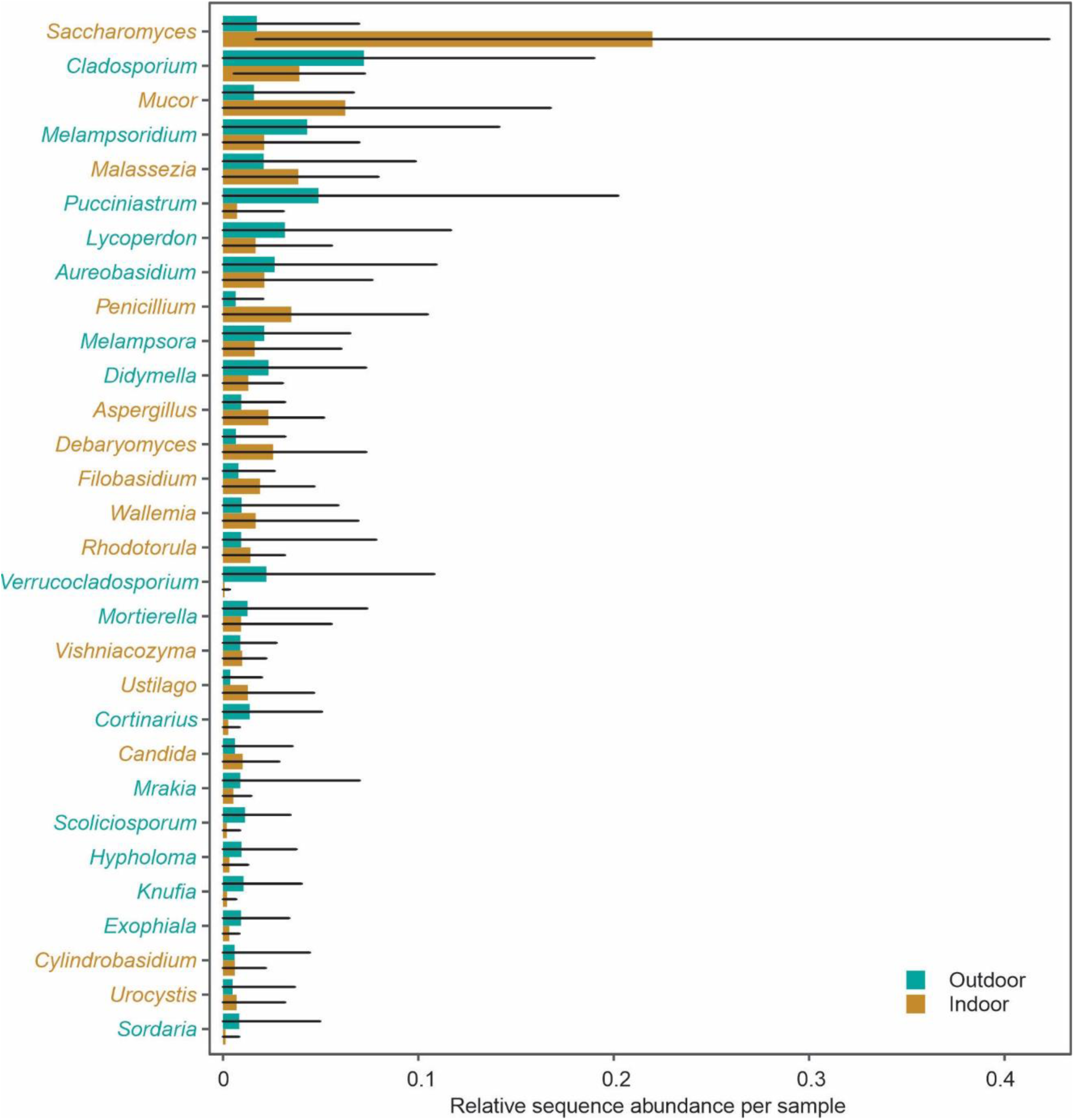
The 30 most abundant genera in the dataset, displaying their average sequence abundance across indoor and outdoor samples in the 125 daycare centers. For the indoor samples, a mean value from the merged bathroom and main room sample was used for the calculations. Genera with higher indoor abundance are displayed in brown color, while genera with higher outdoor abundance are shown in cyan. The black lines indicate standard error.

## Discussion

### Factors influencing the indoor mycobiome

We observed a clear separation between the outdoor and indoor mycobiome across the 125 Norwegian daycare centers, and that numerous variables associated with both the outdoor climate and the indoor environment together influenced the indoor mycobiome. We observed a similar pattern in a recent study of private homes across the same climatic gradients in Norway (24). Likewise, Barberán et al. reported a similar trend from analyzes of house dust mycobiomes throughout the USA (25). However, other preceding studies pointed that indoor air and dust mostly consist of outdoor fungi that have spread into buildings through the ventilation system, windows or doors (12, 30, 31). For example, Shin et al. (2015) concluded that human activity had little influence on the indoor fungal community composition in daycare centers in Seoul, South Korea (31). Similarly, in a study investigating indoor fungi in a housing facility in California, Adams et al. (2013) showed that the outdoor air and not the residents structured the indoor mycobiome (12). Interestingly, in our recent study on seasonality of the indoor mycobiome, the indoor environment was more influenced by the outdoor fungal diversity during summer and fall (10). Thus, as the citizen scientists in the present study sampled during early spring, we may have detected a stronger influence of indoor variables compared to e.g. Shin et al. (2015), where samples were collected from August to October in a comparable climate in South Korea.

According to the CCA analysis, the number of children in daycares accounted for some of the overall variation in the indoor mycobiome composition, together with construction year and the occurrence of pests and rodents. Furthermore, the variables building type, number of departments, room (main room versus bathroom), and type of daycare, correlated significantly with the mycobiome composition in single factor analyses. Taken together, these results indicate that the organization of daycare centers and building features influence the indoor mycobiome composition. In addition to building variables, regional climate related factors such as maximum temperature in June, mean temperature of the coldest quarter and total insolation, also correlated significantly with the indoor mycobiome composition. Longitude, an approximation for regional climate variability, also had explanatory power. Throughout most of Norway, longitude mirrors a climate gradient from oceanic and humid areas in the west, to areas with dryer, colder and high temperature seasonality conditions in the east. Our findings mirror the observations by Barberán et al. (25) and Martin-Sanchez et al. (12), where regional climate was found to be important for the indoor mycobiome. The climate factors most likely have indirect effects on the indoor fungi, as they influence and structure the outdoor fungi that disperse into buildings.

Despite that several of the assessed variables were significantly related to the composition of the indoor mycobiome, only a small fraction of the variation in indoor mycobiome composition was accounted for (7%). However, the low level of explanatory power is not a unique feature to this study, but rather a common trend across most fungal diversity studies (24, 32). Fungal communities are largely assembled through colonization by spore dispersal, which to a large extent is a random process. Because of this, it is generally hard to account for all variables structuring the fungal community composition.

### Taxonomic composition of daycare mycobiome

The most marked taxonomic difference in the indoor and outdoor dust mycobiome was the predominance of yeasts and molds inside the daycare centers. *Saccharomyces* was by far the most abundant genus in our study and had a clear affinity to indoor environments. *Saccharomyces* may partly be derived from food, but has also been found as one of the most abundant genera in the human gut (33) and on children’s skin (34). Other true yeasts, such as *Debaryomyces* and *Candida*, had also a clear affinity to indoor environments in the studied daycares. *Candida* is one of the most widespread genera associated with external (skin) and internal (mouth, digestive tract) parts of the human body (35). It is well documented that *Candida* is particularly associated with children, commonly resulting in oral thrush (massive *Candida* growth in mouth and throat) in the first years of life (36). The lipophilic basidiomycete yeast *Malassezia*, a widespread genus on human skin (18), and *Rhodotorula*, another basidiomycete yeast associated with the human body (35), were also prevalent in the studied daycares. *Malassezia*, as well as *Candida*, are known to be associated with inflammatory skin disorders such as seborrheic dermatitis and atopic dermatitis in childhood as well as in adulthood (37, 38). However, *Malassezia* most often has a commensal role, as they are widespread on healthy skin. For instance, 11 of the 14 known *Malassezia* species were associated with different parts of the skin of 14 healthy adults (20), indicating that human skin is colonized with a wide range of *Malassezia*. On children’s skin, a dominance of the species *Malassezia globosa* has been observed (18). We hypothesize that the yeasts dominating the indoor daycare mycobiome are mainly derived from different parts of the human body. The high density of children and close physical contact may lead to easy and fast transmission of yeasts in daycares, possibly explaining the up-concentration of these species indoors.

In addition to the indoor enrichment of yeasts, several extremotolerant molds, such as *Mucor, Penicillium, Aspergillus* and *Wallemia* also showed a clear preference for the indoor environment. These genera are widespread and prevalent in most of indoor environments (24, 25, 39), and grow rapidly on organic materials. All of these taxa are sometimes detected on and in the human body as well (18). *Cladosporium*, another abundant mold in indoor environments, was prevalent in both indoor and outdoor samples and might largely be dispersed from outdoor sources. Though no direct cause-effect relationship has been established, some of these mold taxa were abundant in houses with children with allergies and respiratory diseases (15, 40).

### Concluding remarks

Taken together, we conclude that the indoor mycobiome of Norwegian daycare centers are dominated by yeasts and molds, and that a multitude of factors structure its composition. For the current study, dust samples were obtained during a relative short time window during the spring 2018. From other studies, we know there is an extensive temporal variability (10), which is not accounted for here. Moreover, sampling at approximately the same time throughout Norway, a country that spans a wide range of latitudes and longitudes, implies that the outdoor climate, vegetation and fungal communities are in different (phenological) growth phases. Such differences can influence which fungi we recovered from different regions we sampled. Indeed, the variable (sampling) month was significantly correlated to the fungal community composition, but it only accounted for a small amount of the variation. Most likely, indoor fungi dominated by yeasts and molds, can be sampled in higher proportions during winter and spring in the Norwegian climate, while the influence of outdoor fungi on the indoor mycobiome will be greater during the growth and sporulation period of most mushrooms (summer and fall). Hence, a sampling time during the winter period may be even more representative of the specific indoor fungal community in future studies.

In the current study, we carried out a citizen science sampling approach for obtaining the material from daycare centers. In line with our previous study (13), only a few outlier samples occurred, and the indoor and outdoor dust samples were largely separated in species composition, indicating a low influence of sampling bias. Moreover, very few samples were discarded due to low DNA yields. Our results supported that citizen science sampling is a powerful approach to obtain samples from a widespread geographic area during a short time span. We advocate for further citizen science studies for evaluating biological and chemical air pollutants, which will also help to raise public awareness on air quality problems in buildings.

## Materials and methods

### Sampling

A list of Norwegian daycare centers was retrieved from the Norwegian Directorate of Health (Helsedirektoratet). The list was sorted alphabetically after counties and municipalities, and the first five municipalities in each county were selected for the study. Likewise, the first 3-4 daycare centers in each of these municipalities were chosen as candidate sites for dust sampling. Sampling kits containing five FLOQSwabs (Copan Italia spa, Brescia, Italy) and a questionnaire were sent to the selected daycare centers with specifications to perform dust sampling on doorframes: (1) outdoor, (2) main room and (3) bathroom. If the daycare had two different departments, we requested to repeat the sampling in (4) the main room and (5) the bathroom of the second department as well. Overall, 572 samples were retrieved from a total of 125 studied daycare centers (Fig. 1), and upon arrival the swabs were stored at -80 °C until DNA extraction.

### DNA extraction and metabarcoding

Samples were prepared and DNA was extracted using the E.Z.N.A Soil DNA kit (Omega Biotek, Norcross, GA, USA). The tips of the swabs were placed in disruptor tubes by using a sterilized scissor. The empty swab tubes were filled with 800 µL SLX-Mlus Buffer to collect the remaining dust before being transferred to the disruptor tubes. The samples were homogenized for 2 × 1 min at 30 Hz using a TissueLyser (Qiagen, Hilden, Germany) and stored at -20 °C until further processing.

DNA extraction and metabarcoding library preparation were performed according to Estensmo et al. (10). Briefly, samples were thawed at 70 °C, followed by an incubation of 10 minutes at the same temperature and homogenized for 2 × 1 min at 30 Hz using the TissueLyser. The samples were then cooled on ice before adding 600 µL chloroform, vortexed and centrifuged at 13 000 rpm for 5 min at RT. The aqueous phase was transferred to a new 1.5 mL tube and an equal volume of XP1 Buffer was added before vortexing. The extract was transferred to the HiBind DNA Mini Column and further processed following the manufacturer’s guidelines. The DNA was eluted in 50 µL Elution Buffer.

We targeted the ITS2 region with the forward primer ITS4: 5′-xCTCCGCTTATTGATATG (41) and the modified reverse primer gITS7: 5′-xGTGARTCATCGARTCTTTG (42), barcodes x ranging from 6-9 base pairs. The amplification mix contained 2 µl DNA template, 14.6 µl Milli-Q water, 2.5 µl 10x Gold buffer, 0.2 µl dNTP’s (25 nM), 1.5 µl reverse and forward primers (10 µM), 2.5 µl MgCl2 (50 mM), 1.0 µl BSA (20 mg/ml) and 0.2 µl AmpliTaq Gold polymerase (5 U/µl). DNA was amplified by initial denaturation at 95 °C for 5 min, followed by 32 cycles of denaturation at 95 °C for 30 s, annealing at 55 °C for 30 s, and elongation at 72 °C for 1 min. A final elongation step was included at 72 °C for 10 min. Amplicons were normalized using the SequalPrep Normalization Plate Kit (Invitrogen, Thermo Fisher Scientific, Waltham, MA, USA) and eluted in 20 μL Elution Buffer. The resulting PCR products were processed into seven libraries of 96 samples using a combination of 96 tagged primers. Each library included 10 technical replicates (dust samples), one mock community (artificial fungal community composed of DNA in 1 ng/µL equimolar concentration from *Mycena belliarum, Pycnoporellus fulgens, Serpula similis* and *Pseudoinonotus dryadeus*), negative DNA controls (using a clean swab as starting material) and negative PCR controls. The 96 PCR products within each library were pooled, concentrated and purified using Agencourt AMPure XP magnetic beads (Beckman Coulter, CA, USA). The quality of the purified pools was measured using Qubit (Invitrogen, Thermo Fisher Scientific, Waltham, MA, USA). The seven libraries were barcoded with Illumina adapters, spiked with PhiX and sequenced in three Illumina MiSeq (Illumina, San Diego, CA, USA) lanes with 2 × 250 bp paired-end reads at Fasteris SA (Plan-les-Ouates, Switzerland).

### Bioinformatics

The bioinformatics analyses were performed according to Estensmo et al. (10). Briefly, raw sequences were demultiplexed independently using CUTADPT (43) allowing no miss-matches between barcode tags and sequence primer, and sequences shorter than 100 bp where discarded. DADA2 (44) was used to filter low quality reads and error correction. We then merged the error corrected sequences using a minimum overlap of five bp. Chimeras were removed using the bimera algorithm, using default parameters implemented in DADA2. The resulting ASV table were further clustered into 10 955 operational taxonomic units (OTUs) using VSEARCH (45) at 97% similarity. LULU (46) was used with default settings to correct for potential OTU over-splitting. Taxonomy was assigned using BLAST (47) to the final OTU table using the UNITE database (48). Sequences with no match to any known fungal sequence and samples with less than 10 OTUs were discarded from downstream analyses. The final raw dataset (without technical replicates and controls) contained 7 399 OTUs and 22 655 516 reads from 572 samples. The number of reads per sample varied from 19 to 182 266 with a mean value of 39 608. The number of OTUs per sample varied from 10 to 863, with a mean value of 257.

### Environmental variables

Metadata about building features and occupancy of each daycare were provided by the volunteers in a questionnaire that were delivered together with the samples (Table 1). The location of daycare centers with complete addresses were provided, and corresponding geographic coordinates (latitude and longitude) were retrieved. Based on these coordinates, relevant environmental variables were extracted and kindly provided by the authors of a recent study modelling the vegetation types in Norway (29). From this extensive set of environmental variables (>30), a subset of non-collinear variables (|r|> 0.6) was selected for further analyses (Table 1).

### Annotation of fungal (OTUs) growth characteristics

We annotated the 1 593 most abundant OTUs, defined as those with >20 sequences and taxonomic annotation at a species, genus or family level, into growth forms/nutritional modes based on literature surveys and information available in the UNITE database (48). Species/genera/families having unknown, dubious or multiple growth forms / nutritional modes, were not included. A complete list of our annotations can be found in Supplementary table 1.

### Statistics

The statistical analyses were performed in R (49).Given that DNA-metabarcoding analyses of samples with low DNA yields may introduce biases during the wet-lab analyses and sequencing, we controlled the consistency of our results. Therefore, the similarity of the technical replicates was evaluated by nonmetric multidimensional scaling (NMDS) using the metaMDS function from the VEGAN package version 2.4-2 (50), and the results were visualized by GGPLOT2 (51) (Fig. S1). As visualized in Fig. S1, the distances between biological replicates are generally markedly higher than between the technical replicates. Then, all the samples in the complete dataset were rarefied to 1 342 sequences using the function rrarefy (VEGAN). Fourteen samples were discarded for downstream statistical analyses due to a lower sequencing depth.

To visualize and investigate patterns in OTU composition in relation to environmental variables, we performed a global non-metric multidimensional scaling (GNMDS) using the VEGAN package and the settings as recommended by the authors. To ensure reliability of the results a detrended correspondence analyses (DCA) was performed in parallel. Extreme outliers common to both ordinations, were manually inspected and subsequently removed from the dataset before the analyses were repeated. Both ordination analyses revealed the same overall pattern (data not shown) and we hereafter focus on the GNMDS analyses. The GNMDS was scaled into half change units and subjected to varimax rotation using principal component analyses (PCA). To confirm convergence the two best solutions of the GNMDS were compared using Procrustes comparisons with 999 permutations (corr: 0.99, p = 0.001). The ordinations were first conducted on the entire dataset containing both indoor and outdoor samples, where a clear pattern was observed. Thereafter, a dataset containing only indoor samples from main rooms and bathrooms was extracted, and the ordinations were conducted on this dataset using the same settings and correlation in the Procrustes comparisons. The following analyses were only conducted on the indoor dataset. The envfit function in VEGAN (i.e. the fit (R^2^) of each variables assessed with a Monte-Carlo analyses of 999 permutation) was used to fit the environmental variables: building type, average construction year, June temperature, longitude, mean temp of the coldest quarter, month, number of departments and children, presence of rodents (pests), proximity to all types of water, proximity to coast, room type, and type of daycare, to the GNMDS. The numerical variables were visualized using the vectors from the output from the envfit function. We further did a variation partitioning with CCA (canonical correspondence analysis) with 999 permutations, to quantify the components of variation by the variables mentioned above, with forward selection, as implemented in vegan.

To investigate OTU-richness trends, a linear mixed effect model was applied using the NLME package (52), including daycare ID as a random contribution. Collinear variables were excluded as described above (|r| > 0.6), however, to further avoid multicollinearity in the mixed effect model the corvif function described in Zuur et al (2009) was applied, using a threshold of 2.5 (53). Backwards stepwise model selection was performed based on Akaike information criterion (AIC). The distribution of the 30 most abundant genera across indoor and outdoor samples were visualized. In addition, bar charts were made based on relative abundances of the rarified OTU table and fungal annotations, comparing the indoor and outdoor environment. To obtain a balanced indoor/outdoor dataset, one indoor “observation” and one outdoor “observation”, we collapsed the indoor samples to one “observation” by using the average values from the indoor samples (main room and bathroom).

### Data accessibility

Our initial dataset, as well as the final rarefied dataset, are available at Dryad together with information about metadata, scripts for bioinformatics and statistics, taxonomic annotations and growth form / nutritional mode annotations.

## Acknowledgements

We would like to acknowledge the daycare centers for the sampling and for providing metadata of the building and occupancy features. Mycoteam AS contributed to the sampling and provided sampling equipment. The research was financially supported by the University of Oslo and the Norwegian Asthma and Allergy Association (NAAF). PMMS was funded by the European Union’s Horizon 2020 research and innovation program (Marie Skłodowska-Curie Individual Fellowship; grant agreement MycoIndoor No 741332).

## References

1. Horner WE. 2003. Assessment of the indoor environment: evaluation of mold growth indoors. Immunology and Allergy Clinics 23:519–531.

2. Nevalainen A, Täubel M, Hyvärinen A. 2015. Indoor fungi: companions and contaminants. Indoor Air 25:125–156.

3. Bornehag CG, Blomquist G, Gyntelberg F, Jarvholm B, Malmberg P, Nordvall L, Nielsen A, Pershagen G, Sundell J. 2001. Dampness in buildings and health. Nordic interdisciplinary review of the scientific evidence on associations between exposure to “dampness” in buildings and health effects (NORDDAMP). Indoor Air 11:72–86.

4. Mendell MJ, Mirer AG, Cheung K, Tong M, Douwes J. 2011. Respiratory and allergic health effects of dampness, mold, and dampness-related agents: A review of the epidemiologic evidence. Environmental Health Perspectives 119:748–756.

5. Portnoy JM, Kwak K, Dowling P, VanOsdol T, Barnes C. 2005. Health effects of indoor fungi. Annals of Allergy, Asthma & Immunology 94:313–320.

6. Madureira J, Paciência I, Rufo J, Ramos E, Barros H, Teixeira JP, de Oliveira Fernandes E. 2015. Indoor air quality in schools and its relationship with children’s respiratory symptoms. Atmospheric Environment 118:145–156.

7. Vandenborght L-E, Enaud R, Urien C, Coron N, Girodet P-O, Ferreira S, Berger P, Delhaes L. 2020. Type 2–high asthma is associated with a specific indoor mycobiome and microbiome. Journal of Allergy and Clinical Immunology doi:https://doi.org/10.1016/j.jaci.2020.08.035.

8. Korkalainen M, Täubel M, Naarala J, Kirjavainen P, Koistinen A, Hyvärinen A, Komulainen H, Viluksela M. 2017. Synergistic proinflammatory interactions of microbial toxins and structural components characteristic to moisture-damaged buildings. Indoor Air 27:13–23.

9. Madureira J, Paciência I, Rufo JC, Pereira C, Teixeira JP, de Oliveira Fernandes E. 2015. Assessment and determinants of airborne bacterial and fungal concentrations in different indoor environments: Homes, child day-care centres, primary schools and elderly care centres. Atmospheric Environment 109:139–146.

10. Estensmo EL, Morgado L, Maurice S, Martin-Sanchez PM, Engh IB, Mattsson J, Kauserud H, Skrede I. 2021. Spatiotemporal variation of the indoor mycobiome in daycare centers. Microbiome doi:10.21203/rs.3.rs-147547/v1.

11. Barberán A, Ladau J, Leff JW, Pollard KS, Menninger HL, Dunn RR, Fierer N. 2015. Continental-scale distributions of dust-associated bacteria and fungi. Proceedings of the National Academy of Sciences 112:5756–61.

12. Adams RI, Miletto M, Taylor JW, Bruns TD. 2013. Dispersal in microbes: fungi in indoor air are dominated by outdoor air and show dispersal limitation at short distances. The ISME Journal 7:1262–1273.

13. Martin-Sanchez PM, Estensmo EF, Morgado LN, Maurice S, Engh IB, Skrede I, Kauserud H. 2021. Analysing indoor mycobiomes through a large-scale citizen science study in Norway. Mol Ecol 30:2689–2705.

14. Prussin AJ, Marr LC. 2015. Sources of airborne microorganisms in the built environment. Microbiome 3:78.

15. Dannemiller KC, Gent JF, Leaderer BP, Peccia J. 2016. Influence of housing characteristics on bacterial and fungal communities in homes of asthmatic children. Indoor air 26:179–192.

16. Ghannoum MA, Jurevic RJ, Mukherjee PK, Cui F, Sikaroodi M, Naqvi A, Gillevet PM. 2010. Characterization of the oral fungal microbiome (mycobiome) in healthy individuals. PLOS Pathogens 6:e1000713.

17. Dupuy AK, David MS, Li L, Heider TN, Peterson JD, Montano EA, Dongari-Bagtzoglou A, Diaz PI, Strausbaugh LD. 2014. Redefining the human oral mycobiome with improved practices in amplicon-based taxonomy: discovery of malassezia as a prominent commensal. PLOS ONE 9:e90899.

18. Jo J-H, Deming C, Kennedy EA, Conlan S, Polley EC, Ng W-I, Segre JA, Kong HH. 2016. Diverse human skin fungal communities in children converge in adulthood. Journal of Investigative Dermatology 136:2356–2363.

19. White TC, Findley K, Dawson TL, Scheynius A, Boekhout T, Cuomo CA, Xu J, Saunders CW. 2014. Fungi on the skin: dermatophytes and Malassezia. Cold Spring Harbor Perspectives in Medicine 4:a019802.

20. Findley K, Oh J, Yang J, Conlan S, Deming C, Meyer JA, Schoenfeld D, Nomicos E, Park M, Kong HH, Segre JA. 2013. Topographic diversity of fungal and bacterial communities in human skin. Nature 498:367–70.

21. Nilsson RH, Anslan S, Bahram M, Wurzbacher C, Baldrian P, Tedersoo L. 2019. Mycobiome diversity: high-throughput sequencing and identification of fungi. Nat Rev Microbiol 17:95–109.

22. Weikl F, Tischer C, Probst AJ, Heinrich J, Markevych I, Jochner S, Pritsch K. 2016. Fungal and bacterial communities in indoor dust follow different environmental determinants. PLOS ONE 11:e0154131–e0154131.

23. Yamamoto N, Hospodsky D, Dannemiller KC, Nazaroff WW, Peccia J. 2015. Indoor emissions as a primary source of airborne allergenic fungal particles in classrooms. Environmental Science & Technology 49:5098–106.

24. Martin-Sanchez PM, Estensmo ELF, Morgado LN, Maurice S, Engh IB, Skrede I, Kauserud H. 2020. Analyzing indoor mycobiomes through a large-scale citizen science study of houses from Norway. Research Square doi:10.21203/rs.3.rs-40337/v1.

25. Barberán A, Dunn RR, Reich BJ, Pacifici K, Laber EB, Menninger HL, Morton JM, Henley JB, Leff JW, Miller SL, Fierer N. 2015. The ecology of microscopic life in household dust. Proceedings of the Royal Society B: Biological Sciences 282:20151139.

26. Gura T. 2013. Citizen science: amateur experts. Nature 496:259–261.

27. Cohn JP. 2008. Citizen science: can volunteers do real research? BioScience 58:192–197.

28. Dickinson JL, Zuckerberg B, Bonter DN. 2010. Citizen science as an ecological research tool: challenges and benefits. Annual Review of Ecology, Evolution, and Systematics 41:149–172.

29. Horvath P, Halvorsen R, Stordal F, Tallaksen LM, Tang H, Bryn A. 2019. Distribution modelling of vegetation types based on area frame survey data. Applied Vegetation Science 22:547–560.

30. Adams RI, Miletto M, Taylor JW, Bruns TD. 2013. The Diversity and Distribution of Fungi on Residential Surfaces. PLOS ONE 8:e78866.

31. Shin S-K, Kim J, Ha S-m, Oh H-S, Chun J, Sohn J, Yi H. 2015. Metagenomic Insights into the Bioaerosols in the Indoor and Outdoor Environments of Childcare Facilities. PLOS ONE 10:e0126960.

32. Peay KG, Kennedy PG, Talbot JM. 2016. Dimensions of biodiversity in the Earth mycobiome. Nature Reviews Microbiology 14:434–447.

33. Raimondi S, Amaretti A, Gozzoli C, Simone M, Righini L, Candeliere F, Brun P, Ardizzoni A, Colombari B, Paulone S, Castagliuolo I, Cavalieri D, Blasi E, Rossi M, Peppoloni S. 2019. Longitudinal Survey of Fungi in the Human Gut: ITS Profiling, Phenotyping, and Colonization. Frontiers in Microbiology 10.

34. Zhu T, Duan Y-Y, Kong F-Q, Galzote C, Quan Z-X. 2020. Dynamics of Skin Mycobiome in Infants. Frontiers in Microbiology 11.

35. Cui L, Morris A, Ghedin E. 2013. The human mycobiome in health and disease. Genome Medicine 5:63.

36. Jean J, Goldberg S, Khare R, Bailey LC, Forrest CB, Hajishengallis E, Koo H. 2018. Retrospective Analysis of Candida-related Conditions in Infancy and Early Childhood Caries. Pediatric dentistry 40:131–135.

37. Nakabayashi A, Sei Y, Guillot J. 2000. Identification of Malassezia species isolated from patients with seborrhoeic dermatitis, atopic dermatitis, pityriasis versicolor and normal subjects. Med Mycol 38:337–41.

38. Zinkeviciene A, Vaiciulioniene N, Baranauskiene I, Kvedariene V, Emuzyte R, Citavicius D. 2011. Cutaneous yeast microflora in patients with atopic dermatitis. Central European Journal of Medicine 6:713.

39. Shelton BG, Kirkland KH, Flanders WD, Morris GK. 2002. Profiles of Airborne Fungi in Buildings and Outdoor Environments in the United States. Applied and Environmental Microbiology 68:1743–1753.

40. O’Connor G T, Walter M, Mitchell H, Kattan M, Morgan WJ, Gruchalla RS, Pongracic JA, Smartt E, Stout JW, Evans R, Crain EF, Burge HA. 2004. Airborne fungi in the homes of children with asthma in low-income urban communities: The Inner-City Asthma Study. J Allergy Clin Immunol 114:599–606.

41. White TJ, Bruns T, Lee S, Taylor J. 1990. Amplification and direct sequencing of fungal ribosomal RNA genes for phylogenetics. PCR Protocols: a Guide to Methods and Applications 18:315–322.

42. Ihrmark K, Bödeker I, Cruz-Martinez K, Friberg H, Kubartova A, Schenck J, Strid Y, Stenlid J, Brandström-Durling M, Clemmensen KE. 2012. New primers to amplify the fungal ITS2 region–evaluation by 454-sequencing of artificial and natural communities. FEMS Microbiology Ecology 82:666–677.

43. Martin M. 2011. Cutadapt removes adapter sequences from high-throughput sequencing reads. EMBnet journal 17:10–12.

44. Callahan BJ, McMurdie PJ, Rosen MJ, Han AW, Johnson AJA, Holmes SP. 2016. DADA2: high-resolution sample inference from Illumina amplicon data. Nature Methods 13:581.

45. Rognes T, Flouri T, Nichols B, Quince C, Mahé F. 2016. VSEARCH: a versatile open source tool for metagenomics. PeerJ Preprints 4:e2409v1.

46. Frøslev TG, Kjøller R, Bruun HH, Ejrnæs R, Brunbjerg AK, Pietroni C, Hansen AJ. 2017. Algorithm for post-clustering curation of DNA amplicon data yields reliable biodiversity estimates. Nature Communications 8:1188.

47. Altschul SF, Gish W, Miller W, Myers EW, Lipman DJ. 1990. Basic local alignment search tool. Journal of Molecular Biology 215:403–410.

48. Koljalg U, Larsson KH, Abarenkov K, Nilsson RH, Alexander IJ, Eberhardt U, Erland S, Hoiland K, Kjoller R, Larsson E, Pennanen T, Sen R, Taylor AF, Tedersoo L, Vralstad T, Ursing BM. 2005. UNITE: a database providing web-based methods for the molecular identification of ectomycorrhizal fungi. New Phytologist 166:1063–8.

49. Team RC. 2019. R: A language and environment for statistical computing. R Foundation for Statistical Computing. Austria: Vienna, https://www.R-project.org/.

50. Oksanen J, Kindt R, Legendre P, O’Hara B. 2019. The vegan package.

51. Wickham H. 2016. ggplot2: elegant graphics for data analysis. springer.

52. Pinheiro J, Bates D, DebRoy S, Sarkar D, Team RC. 2019. nlme: Linear and nonlinear mixed effects models. R package version 3:109.

53. Zuur A, Ieno EN, Walker N, Saveliev AA, Smith GM. 2009. Mixed effects models and extensions in ecology with R. Springer Science & Business Media.

